# Role of the N- and C-terminal regions of FliF, the MS ring component in *Vibrio* flagellar basal body

**DOI:** 10.1101/2021.01.04.425360

**Authors:** Seiji Kojima, Hiroki Kajino, Keiichi Hirano, Yuna Inoue, Hiroyuki Terashima, Michio Homma

**Author notes:** Corresponding author: Division of Biological Science, Graduate School of Science, Nagoya University, Chikusa-ku, Nagoya 464–8602, Japan; E-mail address (to M.H.) and (to S.K.). Institute of Tropical Medicine, Nagasaki University, Nagasaki, 852-8523, Japan (Hiroyuki Terashima). Both authors contributed equally to this work (Seiji Kojima, Hiroki Kajino). Author contributions: S.K. and M.H. designed research; H.K., K.H., Y.I., H.T. and S.K. performed experiments; H.K., S.K., and M.H. analyzed data; H.K., S.K. and M.H. wrote the paper.

## Abstract

The MS ring is a part of the flagellar basal body and formed by 34 subunits of FliF, which consists of a large periplasmic region and two transmembrane segments connected to the N- and C-terminal regions facing the cytoplasm. A cytoplasmic protein, FlhF, which determines the position and number of the basal body, supports MS ring formation in the membrane. In this study, we constructed FliF deletion mutants that lack 30 or 50 residues at the N-terminus (ΔN30 and ΔN50), and 83 (ΔC83) or 110 residues (ΔC110) at the C-terminus. The N-terminal deletions were functional and conferred motility of *Vibrio* cells, whereas the C-terminal deletions were nonfunctional. The mutants were expressed in *Escherichia coli* to determine whether an MS ring could still be assembled. When co-expressing ΔN30FliF or ΔN50FliF with FlhF, fewer MS rings were observed than with the expression of wild-type FliF, in the MS ring fraction, suggesting that the N-terminus interacts with FlhF. MS ring formation is probably inefficient without an additional factor or FlhF. The deletion of the C-terminal cytoplasmic region did not affect the ability of FliF to form an MS ring because a similar number of MS rings were observed for ΔC83FliF as with wild-type FliF, although further deletion of the second transmembrane segment (ΔC110FliF) abolished it. These results suggest that the terminal regions of FliF have distinct roles; the N-terminal region for efficient MS ring formation and the C-terminal region for MS ring function. The second transmembrane segment is indispensable for MS ring assembly.

**Importance:** The bacterial flagellum is a supramolecular architecture involved in cell motility. At the base of the flagella, a rotary motor that begins to construct an MS ring in the cytoplasmic membrane comprises 34 transmembrane proteins (FliF). Here, we investigated the roles of the N and C terminal regions of FliF, which are MS rings. Unexpectedly, the cytoplasmic regions of FliF are not indispensable for the formation of the MS ring, but the N-terminus appears to assist in ring formation through recruitment of FlhF, which is essential for flagellar formation. The C-terminus is essential for motor formation or function.

## Introduction

Bacteria are prokaryotes, approximately 1 µm in length. They travel in fluids using flagella that extend from the cell surface. The flagella are assembled as a supramolecular structure composed of more than 20 types of component proteins (1–3). A rotary motor at the base of each flagellum serves as a power engine. The motor uses the electrochemical potential difference of the coupling ions across the cell membrane to generate a rotational force. The cells can move by rotating helical flagellar filaments as screws. Bacteria use different coupling ions. *Escherichia coli* and *Salmonella enterica* have H^+^-driven motors, or *Vibrio alginolyticus* and alkalophilic *Bacillus* have Na^+^-driven motors (4, 5). The flagellar motor is composed of a stator and rotor, and a dozen stator units are assembled around each rotor (6). Structural changes in the stator, that couple with the flow of ions, generate torque by interacting with the rotor (5, 7). Bacteria containing Na^+^-driven flagella, such as marine *Vibrio*, have two transmembrane proteins, PomA and PomB, as stator proteins, and form a heteromultimer complex (8, 9).

The rotor consists of an MS ring located on the cell membrane, and a C ring built on the cytoplasmic side of the MS ring. The MS ring is constructed by assembling dozen copies of FliF, a protein with two transmembrane segments (10–12). The subatomic structure of the MS ring was determined by cryo-electron microscopy, although the N-terminal and C-terminal regions were not found in *S. enterica* (13, 14). The C ring is composed of three proteins, FliG, FliM, and FliN, and forms a complex with the MS ring via FliG (15). FliG plays an important role in the generation of rotational force, which is generated by the interaction between the stator protein MotA and the rotor protein FliG in *E. coli* (16, 17). The interaction to generate torque similarly occurs in the sodium-driven motor of *V. alginolyticus* (18–20).

Bacteria form flagella with various numbers and positions depending on the species. *E. coli* and *S. enterica* cells have peritrichous flagella with approximately 8–10 flagella per cell, which grow randomly on the cell surface. Contrarily, *Pseudomonas aeruginosa* and *Vibrio* species have a single flagellum at one pole with the cytoplasmic proteins FlhF and FlhG determining their position and number (21–24). FlhF, which has a GTPase activity and is similar to the signal recognition particle protein Ffh which has a role in protein export, controls the number of flagella positively localized at the cell pole, to determine the position of the flagellum. On the other hand, FlhG, which has ATPase activity and is similar to the cell division inhibitor MinD, which represses FlhF function to negatively control the number of flagella. In addition to these two proteins, as for *V. alginolyticus*, it has been shown that HubP and SflA is involved in the polar flagellar formation (24-26).

It is known that FlhG acts on FlhF to negatively control flagellar formation (22). The interaction between FlhF and FlhG has been shown by a pull-down assay, and the polar localization of FlhF increases in the absence of FlhG (27). Although it has not been shown in *V. alginolyticus*, FlhG (named FleN in *P. aeruginosa*) in *V. cholerae* (23) and in *P. aeruginosa* (28) interacts with the master transcription factor of flagellar genes, FlrA (*V. cholerae*) or FleQ (*P. aeruginosa*) to negatively regulate transcription.

The flagellum is constructed by sequentially assembling the flagellar structural proteins on the MS ring (29). The extracellular axial structures are called rods, hooks, and filaments in close proximity to the MS ring. The component proteins were supplied by the export apparatus, which is located inside the MS ring. It has been shown that FlhA, which is one of the main components of the export apparatus, interacts with FliF (30). We speculated that the formations of the MS ring and export apparatus are dependent on each other. In *E. coli*, FlhA and FliG are required for MS ring formation to assemble FliF (31). Contrarily, it has been reported that *Salmonella* MS ring requires FliG but not FlhA to assemble FliF (32). Furthermore, it has been shown that *Salmonella* FliF can form MS rings by its overexpression alone (33, 34). In the case of *Vibrio* FliF, the overproduction of FliF in *E. coli* forms a small amount of MS ring, and co-expression of FliG or FlhF promotes MS ring formation (35).

In this study, we aimed to clarify how the cytoplasmic protein FlhF, which positively controls the number of flagella, acts on the MS ring component protein FliF to promote MS ring formation. We examined whether the cytoplasmic N-terminal or C-terminal region of FliF, which is likely to interact with FlhF, affects MS ring formation.

## Results

### Motility of *fliF* mutants lacking the N-terminal and C-terminal regions

The *V. alginolyticus* FliF protein is composed of 580 amino acids with two transmembrane (TM) segments, and its N-terminal 54 residues and C-terminal 87 residues facing the cytoplasm (Fig. 1A). We constructed deletion mutants removing 30 or 50 N-terminal residues (ΔN30 and ΔN50 respectively), and 83 or 110 C-terminal residues (ΔC83 and ΔC110 respectively). (Fig. 1B, Fig. S1). The resulting mutant proteins were expressed from the plasmid in *Vibrio fliF*-deficient strains (NMB196) and examined for cell motility on a soft agar plate (Fig. 1C). The N-terminal deletion mutants were able to form a swimming ring, although the ΔN50 ring was smaller than that formed by the wild type. The amount of FliF protein was reduced in the ΔN50 mutant, suggesting that the deletion mutant was unstable or expressed in lower amounts (Fig. S2). The flagellar formation and the swimming speed were similar among the mutants, wen examined in a high-intensity dark-field microscope. The C-terminal deletions abolished the motility of the cells on the soft agar plate (Fig. 1C). No flagella formation was observed in these mutants in a high-intensity dark-field microscope.

**Fig. 1.**
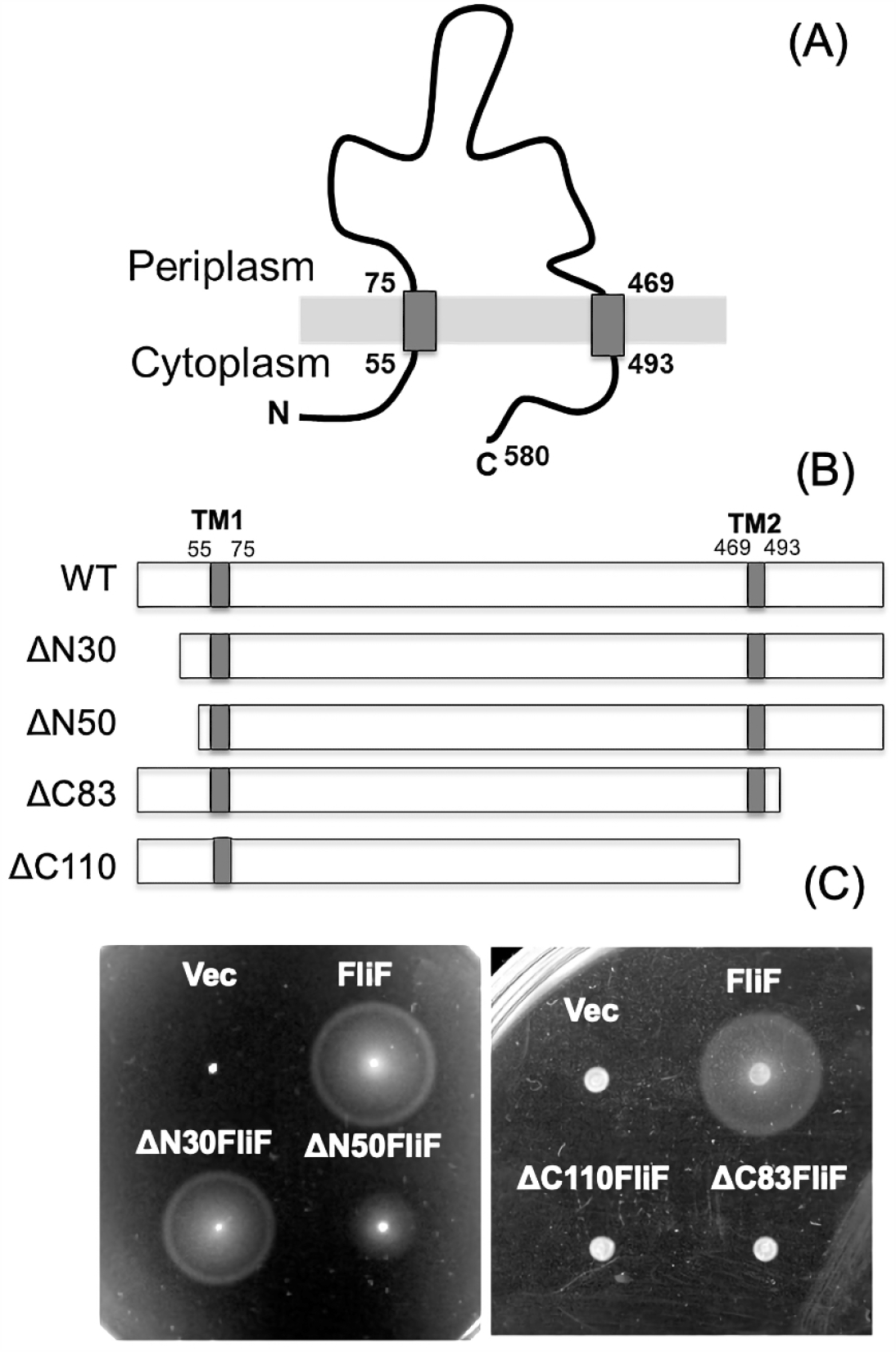
(A) Membrane topology of *Vibrio* FliF. (B) Schematic diagram of N-terminal and C-terminal deleted *Vibrio* FliF. (C) Motility of N-terminal deletion FliF in soft agar plate. The *fliF* deletion mutant (NMB196) producing wild-type FliF, ΔN30FliF, and ΔN50FliF from the pBAD plasmids (Vec) were inoculated on soft agar plate at 30 °C for 5 hours. Motility of C-terminal deletion FliF in soft agar plate. The *fliF* deletion mutant (NMB196) producing wild-type FliF, ΔC83FliF, and ΔC110FliF were inoculated on soft agar plate at 30 °C for 4 h.

### MS ring formation of N-terminal deletions of FliF

We have previously reported that the MS ring is formed by *Vibrio* FliF with the co-expression of FlhF or FliG in *E. coli* cells (35). We cloned ΔN30 or ΔN50 mutant *fliF* into a pCold expression vector with a His-tag and a Factor Xa protease cleavage sequence at the N-terminus. We confirmed that the N-terminal His-tag did not affect FliF function in *Vibrio* cells (Fig. S3). Both mutants were also expressed similarly as the wild-type FliF in *E. coli*, and the membrane fraction was recovered and solubilized with the detergent dodecyl maltoside (DDM). The FliF protein was purified by Ni-affinity resin chromatography using the fused FliF tag. The affinity-purified fraction was precipitated by ultracentrifugation and used as the MS ring fraction (Fig. S4). MS rings were observed in the MS ring fraction with an electron microscope, for both ΔN30FliF and ΔN50FliF mutants, although in much less amounts than in the wild-type FliF (Fig. 2C and 2E). When FlhF was co-expressed with FliF, MS ring formation was facilitated in wild-type FliF, as reported previously (Fig. 2B) (35), whereas FlhF did not promote MS ring formation in ΔN30FliF or ΔN50FliF mutants (Fig. 2D and 2F). These results suggest that the MS ring formed by the deletion proteins is unstable, and it is difficult to form an MS ring with the assistance of FlhF in *E. coli* cells.

**Fig. 2.**
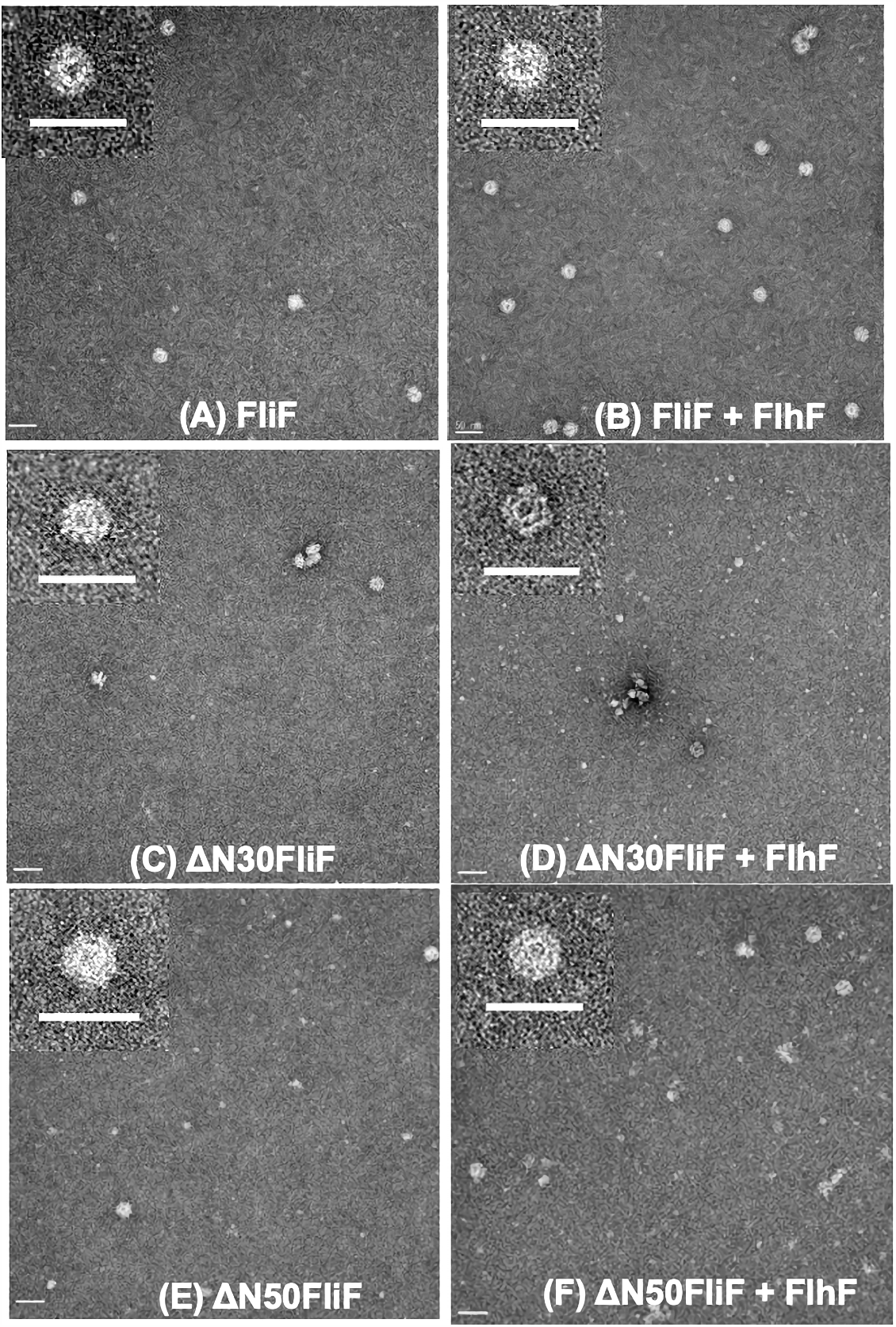
Electron microscopic observation of MS ring made by N-terminal deleted FliF. *E. coli* BL21 (DE3) cells harboring pRO101 (A), pRO101 and pTSK22 (B), pRO101-*ΔN30* (C), pRO101-*ΔN30* and pTSK122 (D), pRO101-*ΔN50* (E), or pRO101-*ΔN50* and pTSK122 (F) were cultured and the MS ring was isolated. The membrane fraction was solubilized with dodecyl maltoside (DDM) and the MS ring was precipitated by ultracentrifugation (Fig. S4). The MS ring fraction was observed with an electron microscope. The scale bars; 50 nm.

### Observation of polar localization of N-terminally deleted FliF

It has been shown that polar FliF localization depends on FlhF, which is localized at the cell poles (35). N-terminal deletion mutants were fused with GFP to construct ΔN30FliF-GFP or ΔN50FliF-GFP, and expressed by the arabinose-inducible plasmid in the *fliF* deletion strain. Fluorescent dots were observed at the cell poles in ΔN30FliF-GFP or ΔN50FliF-GFP, although the localization appeared to be reduced as compared to the wild-type FliF-GFP (Fig.3A). In the *flhF* deletion strain, neither ΔN30FliF-GFP nor ΔN50FliF -GFP showed localization at the poles, as observed for the wild-type FliF-GFP, and fluorescence was observed throughout the cells (Fig. 3B). The *rpoN* mutant does not express polar flagellar genes, except for the master regulator, *flaK*. When ΔN30FliF-GFP, ΔN50FliF-GFP, or FliF-GFP were co-expressed with FlhF in the *rpoN* mutant, the fluorescence dots were observed at the cell poles (Fig. 3C), indicating that ΔN30FliF and ΔN50FliF can penetrate the cell pole independently. The above results supported the idea that the N-terminal cytoplasmic region of FliF preceding the transmembrane segment was not an interaction site for FlhF, but it has a role in promoting MS ring formation.

**Fig. 3.**
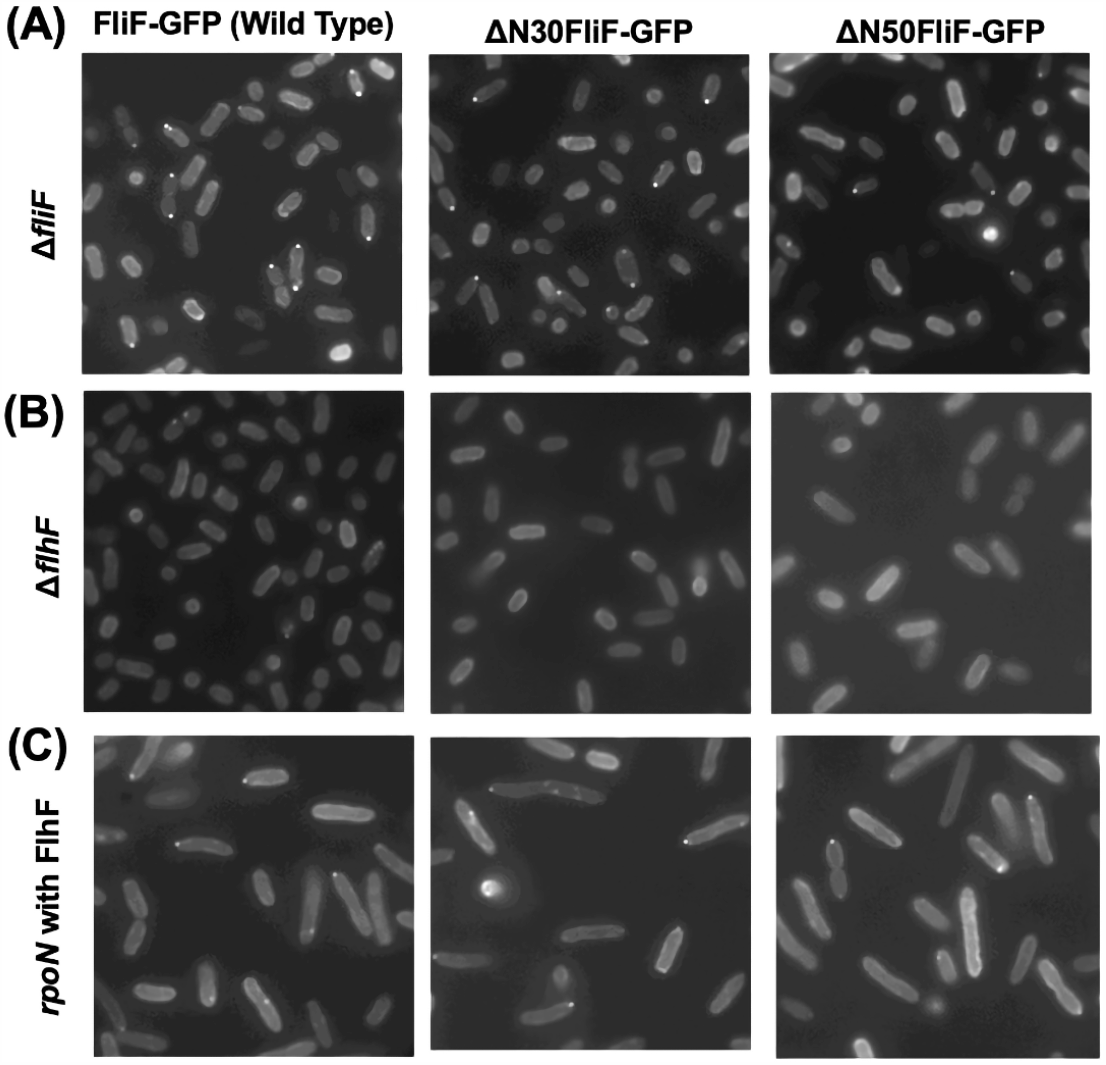
Observation of localization of N-terminal deleted FliF in *Vibrio*. The cells of Δ*fliF* strain (A), Δ*flhF* strain (B), or *flhF* co-expressed *rpoN* strain (C), harboring pYI101(FliF-GFP), pYI101-*ΔN30* (ΔN30FliF-GFP), pYI101-*ΔN50* (ΔN50FliF-GFP) were cultured in VPG broth containing 0.02% (w/v) arabinose for 4 h at 30 °C and were observed by fluorescent microscopy.

### The ability of C-terminally deleted FliF to form an MS ring

The C-terminal deletion mutants were overexpressed in *E. coli* in the same manner as the N-terminal deletion mutants, and an MS ring fraction was obtained (Fig. S4). When the MS ring fraction was observed under an electron microscope, it was similar for ΔC83FliF as well as the wild-type FliF (Fig. 4A), and FlhF promoted MS-ring formation (Fig. 4B). Contrarily, further deletion of the second TM segment abolished MS ring formation, as observed by electron microscopy (Fig. 4C).

**Fig. 4.**
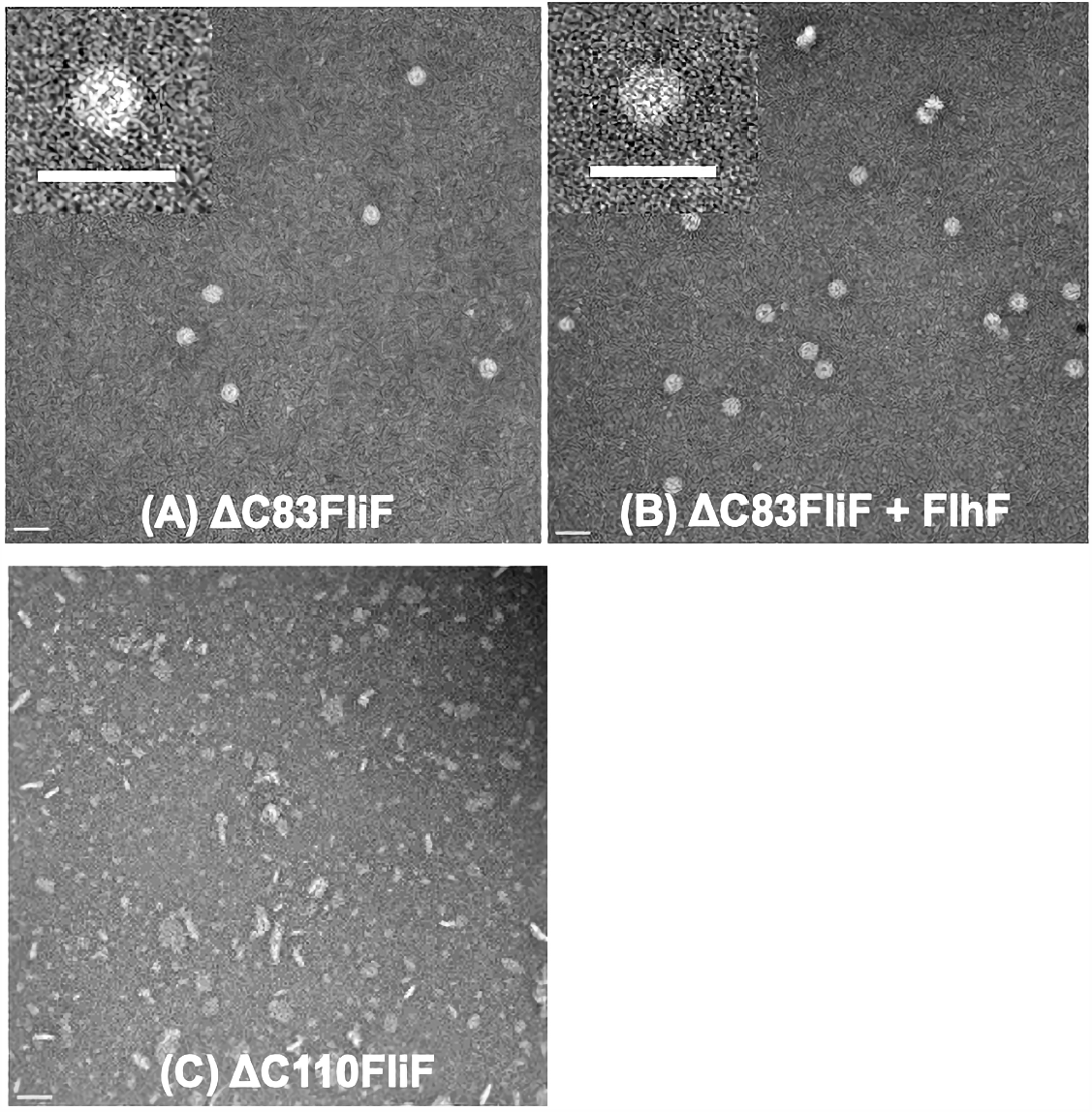
Electron microscopic observation of MS ring made by C-terminal deleted FliF. *E. coli* BL21 (DE3) cells harboring pRO101-*ΔC83* (A), pRO101-*ΔC83* and pTSK22 (B), pRO101-*ΔC110* (C) were cultured and purified same as Fig. 2. The MS ring fraction was observed with an electron microscope. The scale bars; 50 nm.

## Discussion

Bacterial flagella are supramolecular structures composed of tens of thousands of molecules composed of 20 or more components. It is believed that flagella formation begins from the rotor MS ring and C ring assembly, and the export apparatus is constructed beneath the MS ring. After the basal structure has been constructed, the flagellar axial proteins are transported extracellularly by the export apparatus through the interior space of the tubular flagellar structure. Flagellar proteins are assembled at the distal end of the filament to form a tubular structure. To construct a flagellar structure, it is essential to form an MS ring as a starting point. It has been shown that *Salmonella* FliF alone can form an MS ring if the protein is overproduced (33), whereas *Vibrio* FliF requires FliG or FlhF to make the MS ring efficiently (35).

When the *Vibrio* FliF protein was overexpressed in *E. coli*, more than half of the protein was recovered as a soluble protein in the cytoplasmic fraction (36). It was eluted as a broad peak in gel filtration chromatography, with an estimated molecular weight of approximately 700 kDa. The structure seemed to be a multimer composed of approximately 10 FliF molecules. Since this structure could interact with FliG (36), it was presumed that the TM regions were woven inside the structure so that the C-terminal regions were exposed to the outside of the structure. This would be a consequence of many proteins not being inserted into the membrane. In *Vibrio* membrane proteins such as PomA and PomB, most proteins are inserted into the membrane by overexpression (37).

FliF of various species is a protein consisting of from 500 to 600 amino acids and contains two TMs at both ends. Both the N-terminus and C-terminus are present in the cytoplasmic region (Fig. 1A). The C-terminal region is known to interact with the N-terminal region of the FliG protein, which is a C-ring component protein (36). In the extracellular periplasmic region, three ring-building motifs (RBMs) have been inferred from homology with Type III injectisomes (38). These are RBM1, RBM2, and RBM3 beginning from the N-terminus. Recently, structural analysis of the MS ring of *Salmonella* was performed at atomic resolution by single particle analysis using cryo-electron microscopy (13, 14). RBM1, RBM2, and RBM3 contribute to the M ring and S ring formation, respectively. The MS ring formed a ring structure through the interaction of 34 FliF molecules in the extracellular domain. Although the MS ring structure of *V. alginolyticus* has not been solved, it is presumed that it has a similar structure based on its homology and ring size (35).

The cytoplasmic N-terminus of FliF is relatively highly conserved among *Vibrio* species and is approximately 30 amino acids longer than that of the *Salmonella* or *E. coli* FliF. We speculated that this long N-terminal region would prevent MS ring formation in *E. coli*. In addition, since the N-terminal cytoplasmic region of *Vibrio* species has a similar length and homology, it has been speculated that this N-terminal region has a specific function in *Vibrio* FliF. The C-terminal deletion mutants, ΔC83FliF and ΔC110, which lacked 83 and 110 C-terminal residues, completely lost their function, whereas the N-terminal deletion mutants, ΔN30FliF or ΔN50FliF, which lacked 30 and 50 N-terminal residues, were unexpectedly functional in *Vibrio*. The polar localization of the N-terminal deletion mutants was investigated using FliF-GFP. FliF-GFP dots were observed at the poles, although the polar localization was slightly lower than that of the wild type. These polar localizations disappeared as observed in the FlhF-deficient background, suggesting that ΔN30FliF and ΔN50FliF require FlhF to localize at the cell pole. However, FlhF does not promote MS ring formation by using ΔN30FliF or ΔN50FliF in *E. coli* cells. It seems that FlhF can recruit these N-terminally deleted FliF to the pole, but these constructs are unstable to form the MS ring efficiently. Contrarily, we showed that MS rings were formed in the C-terminal deletion ΔC83FliF, and FlhF promoted ring formation using this construct in a manner similar to that of wild-type FliF. The cytoplasmic regions were not essential for the formation of the MS ring. We speculate that the N-terminal cytoplasmic region of FliF is involved in the stability of the MS ring or in FlhF assistance for MS ring formation. Contrarily, the C-terminal cytoplasmic region is not involved in MS ring formation and does not interact with FlhF. Although the formation of the MS ring seems to be normal in the C-terminal cytoplasmic deletion mutant, motility is compromised and the flagellum is not generated. This is because the C-terminal region of FliF interacts with the N-terminal region of FliG to form a C ring. In the C-terminal deletion mutant ΔC110FliF, we could not observe any MS ring, suggesting that at least the second TM region is essential for the formation of the MS ring. As a result, we concluded that the N-terminal region is involved in MS ring formation, although it is not essential, and this region does not interfere with ring formation in *E. coli*. It is not known whether FlhF interacts directly or indirectly with FliF. However, FlhF likely interacts with the N-terminal sequence of FliF. The ring-forming ability and polar localization ability of FliF may not be coordinated with each other. It has been speculated that the assembly of the MS ring requires a core structure, which is a part of the rod composed of FliQ, FliP, and FliQ, is fitted inside the MS ring and consists of a Type III export apparatus (39, 40). FliQ, FliP, and FliQ are 4-fold (25 kDa), 2-fold (9 kDa), and 6-fold (26 kDa) transmembrane proteins respectively, which form a 5:4:1 stoichiometric structure. Presumably, the FliOPQ core is required for FliF assembly to form an MS ring under normal conditions, which does not overproduce the FliF protein.

To form the MS ring, FliF must first be inserted into the membrane. A general membrane transport mechanism is thought to be used for the membrane insertion of the FliF protein (41, 42). First, *fliF* mRNA is recognized by the ribosome, followed by translation. When the TM1 region of FliF is translated, this hydrophobic region is recognized by the SRP protein (Ffh) and interacts with the FtsY membrane-bound to the transport device to target it. FtsY and Ffh are GTPases with a three-domain structure. These proteins are homologous with the GTPase FlhF (27, 35). We propose a scheme to form an MS ring from FliF monomers. Considering the function of FlhF in relation to SRP, FlhF may interact with SRP to facilitate its targeting to the Sec translocon (Fig. 5). In *Vibrio*, SflA prevents flagellar formation in the absence of FlhF because we have shown that the *flhF* and *flhG* double deletion strain, whose cells almost lose flagella, recover the flagellar formation at the peritrichous position by the additional mutation of *sflA*. In the next step, we want to show evidence that SRP interacts with flagellar proteins, such as SflA and FlhF.

**Fig. 5.**
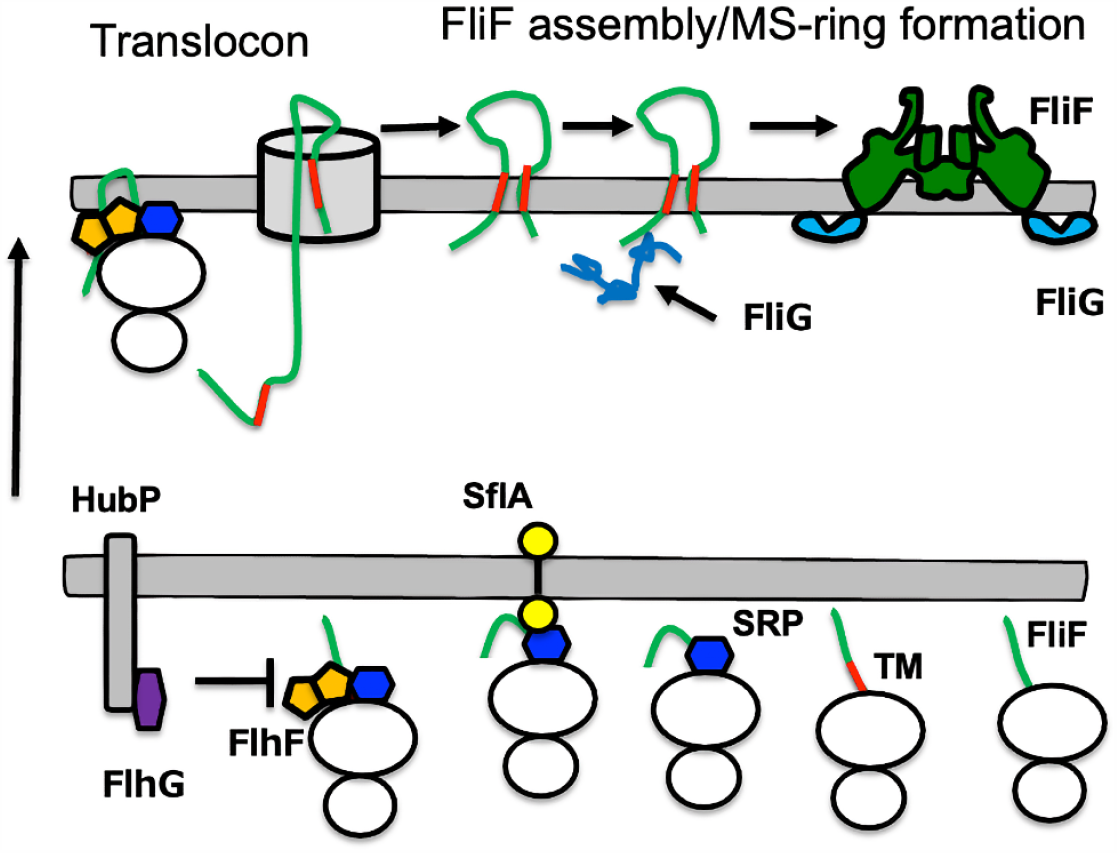
A model of MS ring formation made by FliF proteins. SflA may interact with SRP to prevent FliF to interact with SPR recognition particle. FlhF dominates the SflA protection. FlhF may assist FliF to insert the Sec translocon machinery.

## Materials and methods

### Bacterial strains and plasmids

The bacterial strains and plasmids used in this study are listed in Table S1. *Vibrio* was cultured in VC broth (0.5% [w/v] hipolypeptone, 0.5% (w/v) yeast extract, 3% (w/v) NaCl, 0.4% (w/v) K_2_HPO_4,_ 0.2% (w/v) glucose] or VPG broth [1% (w/v) hipolypeptone, 3% (w/v) NaCl, 0.4% (w/v) K_2_HPO_4,_ 0.5% (w/v) glycerol], and *E. coli* was cultured in LB broth [1% (w/v) bactotryptone, 0.5% (w/v) yeast extract, 0.5% (w/v) NaCl] or SB broth [1.2%(w/v) bactotryptone, 2.4%(w/v) yeast extract, 1.25% (w/v) K_2_HPO_4_, 0.38% (w/v) KH_2_PO_4_, 0.5% (v/v) glycerol]. Chloramphenicol was added to a final concentration of 2.5 µg/mL for *Vibrio* and 25 µg/mL for *E. coli*. Ampicillin was added at a final concentration of 100 µg/mL for *E. coli*. Kanamycin was added at a final concentration of 100 µg/mL for *Vibrio* spp.

### Construction of the deletion mutants

To generate N-terminal FliF deletion constructs, a one-step PCR-based method was employed as previously described (43). To generate C-terminal FliF deletion constructs, a stop codon was introduced at the desired position of *fliF* by the QuikChange site-directed mutagenesis method as described by Stratagene (36). Each mutation was confirmed by DNA sequencing.

### Transformation of *V. alginolyticus*

Introduction of the plasmid into *V. alginolyticus* was performed according to an electroporation method using Gene Pulser (Bio Rad) as previously described (44).

### MS ring purification

*E. coli* BL21(DE3) cells harboring pRO101 or its derivatives (for expression of FliF or its deletions), and pTSK122 (for expression of FlhF) were inoculated from a frozen stock onto a plate containing appropriate antibiotics, and the colonies were inoculated into 30 mL of LB broth and cultured with shaking at 37 °C overnight. 20 mL of the overnight culture was added to 2 L of LB broth and cultured at 37 °C with shaking until OD_600_ = ca. 0.5. The cells were then subjected to cold shock by placing them in ice-cold water for 40 min. After IPTG was added to a final concentration of 0.5 mM, and cultured with shaking at 16 °C overnight. The cells were collected (4,600 × *g*, 10 min) and suspended in TEN buffer (10 mM Tris-HCl [pH 8.0], 5 mM EDTA-NaOH [pH 8.0], 50 mM NaCl) containing proteinase inhibitor (cOmplete, Sigma-Aldrich Co.). The cell suspension was transferred to a 50 mL Falcon tube and stored at −80 °C.

The bacterial suspension was thawed in water, and the cells were disrupted using a French press (9,000–10,000 psi, OTAKE Works). Undisrupted cells were removed by centrifugation (20,000 × *g*, 20 min), and the supernatant was ultracentrifuged at 90,000 × *g* for 1 h. The precipitate was suspended in 45 mL of suspension buffer (10 mM Tris-HCl [pH 8.0], 50 mM NaCl), and 5 mL of 10% dodecyl maltoside (DDM) was added. After stirring at 4 °C for 1 h, the suspension was centrifuged at 20,000 × *g* for 20 min, and the supernatant was ultracentrifuged at 90,000 × *g* for 1 h. The precipitate was suspended in 10 mL of re-suspension buffer (10 mM Tris-HCl [pH 8.0], 5 mM EDTA-NaOH [pH 8.0], and 50 mM NaCl, 0.05% [w/v] DDM) to obtain a crude MS ring fraction. This fraction was shaken in a cold room overnight and centrifuged at 20,000 × *g* for 5 min. The supernatant was used for Ni-affinity purification using a His-tag.

### Ni affinity purification of the MS ring using His-tag

A 10 mL empty column (Qiagen) was packed with 1 mL of Ni-NTA superflow (Qiagen). After washing with 10 mL of MilliQ water, the column was equilibrated with 10 mL of wash buffer (10 mM Tris-HCl [pH 8.0], 5 mM EDTA-NaOH [pH 8.0], 50 mM NaCl, 0.05% [w/v] DDM, 50 mM imidazole). The crude MS ring fraction was added to the column, and the flow-through fraction collected to increase the recovery of the MS ring was added to the column again. The column was washed with 5 mL of wash buffer and again with 20 mL of wash buffer. Thereafter, the protein was eluted with 5 mL of elution buffer (10 mM Tris-HCl [pH 8.0], 5 mM EDTA-NaOH [pH 8.0], 50 mM NaCl, 0.05% [w/v] DDM, and 300 mM imidazole) to obtain the MS ring fraction. The fraction was ultracentrifuged at 90,000 × *g* for 1 h. The precipitate was resuspended in 100 μL of the remaining elution buffer.

### Observation by electron microscopy

The purified MS ring was observed by negative staining using an electron microscope. After hydrophilizing the carbon-coated copper grids, 2.5 µL of the sample solution was placed on the grid and stained with 2% (w/v) uranyl acetate. The grid was observed using a transmission electron microscope (JEM-1010, JEOL) at 100 kV.

### Fluorescence microscopy observations

*Vibrio* cells were cultured overnight in VC medium at 30 °C. The overnight culture was diluted 1:100 in fresh VPG medium containing 0.02% (w/v) arabinose and 100 µg mL^−1^ kanamycin, and was cultured at 30 °C for 4 h.

Fluorescence microscopy was performed as previously described (27). Briefly, cultured cells were harvested and resuspended in V buffer (50 mM Tris-HCl [pH 7.5], 300 mM NaCl, and 5 mM MgCl_2_). These cells were fixed on slides via poly-_L_-lysine, washed with V buffer, and observed under a BX-50 microscope (Olympus). Fluorescent images were recorded and processed using a digital camera (Hamamatsu photonics ORCA-Flash4.0) and HSR imaging software (Hamamatsu Photonics).

## Acknowledgements

We thank Dr. Kimika Maki for technical support with electron microscopy. This work was supported in part by JSPS KAKENHI Grant Numbers 16H04774 (to S.K.), or 20H03220 (to M.H.).

## Supporting information

Supplementary information associated with this article can be found in the online version of the publisher’s website.

